# Albumin-binding Aptamer Chimeras for Improved siRNA Bioavailability

**DOI:** 10.1101/2021.04.15.440012

**Authors:** Jonah C. Rosch, Ella N. Hoogenboezem, Alexander G. Sorets, Craig L. Duvall, Ethan S. Lippmann

## Abstract

Short interfering RNAs (siRNAs) are potent nucleic acid-based drugs designed to target disease driving genes that may otherwise be undruggable with small molecules. However, the potential of administering therapeutic siRNA *in vivo* is limited by poor pharmacokinetic properties, including rapid renal clearance and nuclease degradation. Nanocarriers have traditionally been explored as means to overcome these challenges, but they have intrinsic downsides such as dose-limiting toxicity and synthetic complexity. Backpacking on natural carriers such as albumin, which is present at high concentration and has a long half-life in serum, is an effective way to modify pharmacokinetics of biologic drugs that otherwise have poor bioavailability. In this work, we sought to develop albumin-binding aptamer-siRNA chimeras to improve the bioavailability of siRNA. We used a Systematic Evolution of Ligands through Exponential Enrichment (SELEX) approach to obtain RNA aptamers with modified bases that bind albumin with high affinity. We then fused the aptamers directly to an siRNA to generate the chimera structure. These aptamer-siRNA chimeras are stable in serum, exhibit potent gene knockdown capabilities *in vitro*, and display extended circulation time *in vivo*. We suggest that this albumin-binding aptamer-siRNA chimera approach is a promising strategy for drug delivery applications.

## Introduction

Short interfering RNA (siRNA) is a powerful technology platform for silencing genes that are traditionally undruggable targets by small molecule drugs. By the endogenous interference mechanism, long, double-stranded RNA is cleaved by the endoribonuclease, Dicer, into small siRNA fragments (20-25 bp). The antisense strand of the siRNA is incorporated into the RNA-induced silencing complex (RISC), which recognizes and degrades the complementary mRNA, inhibiting translation of the encoded protein^1, 2^. Despite the potential of siRNA as a therapeutic platform, the challenges of systemic delivery of siRNA have limited its clinical use. siRNA drugs are limited by their rapid degradation and renal clearance^3–5^. Nanocarriers, such as polymeric capsules and liposomes, have been explored as a means to address these challenges, but these drug delivery vehicles often involve complex syntheses and can induce toxicity^3, 6, 7^. We previously reported the utility of direct conjugation of siRNA to a diacyl lipid moiety (siRNA-L_2_) that binds to endogenous serum albumin^8^. Albumin’s extraordinary circulation halflife of 19 days and high concentration in blood (40 mg/mL) make it an ideal target to improve the bioavailability of exogenous drugs^9, 10^. By utilizing albumin as a carrier, the circulation half-life and bioavailability of siRNA were increased, while renal accumulation was reduced. However, the affinity of L_2_ for albumin was moderate (K_d_ of ~1 μM), and its lipid-based structure may lend to nonspecific interactions with other serum components. Hence, we sought an alternative route for targeting albumin using a fully nucleic acid-based system comprising an siRNA fused with an albumin-binding aptamer.

Aptamers are single stranded DNA or RNA molecules that can be selected for binding to a specific target through a process known as Systematic Evolution of Ligands through Exponential Enrichment (SELEX)^11–14^. Direct fusions of siRNA and RNA aptamers, which produces an aptamer-siRNA “chimera” structure, have been previously reported to confer targeting ability to siRNA and siRNA-carriers *in vivo*^15–20^. Though these platforms benefit from the ability of the aptamer to bind target cells with high affinity and specificity, they are still subject to the poor circulation time and rapid clearance of the RNA-based therapeutic^21, 22^. Attempts to overcome this barrier through conjugation to polyethylene glycol (PEG), a hydrophilic polymer known for its ability to increase systemic circulation time of nanoformulations, have been halted due to patient antibody-mediated immune responses^23, 24^. As such, the ability of albumin to be reabsorbed in the kidneys is particularly appealing as a strategy for conferring increased circulation half-life^25^. In this work, we generated RNA aptamers with high affinity for both mouse and human serum albumin. We present *in vitro* and *in vivo* characterization of two aptamer-siRNA chimeras that exhibit enhanced half-life in serum without reductions in gene silencing potency.

## Materials and Methods

### Materials

Human serum albumin (A1653) and mouse serum albumin (A3139) were purchased from Sigma (St Louis, MO). Fluorescently labeled human serum albumin (HS1-S5-1) was purchased from NANOCS (New York, NY). The starting DNA library, primers, and custom aptamer sequences were purchased from Integrated DNA Technologies (San Jose, CA) with HPLC purification. The starting single-stranded DNA (ssDNA) library was synthesized with 40 nucleotide random bases flanked by 20 nucleotide primer ends required to perform PCR amplification (forward fixed region: TCGCACATTCCGCTTCTACC, reverse fixed region: CGTAAGTCCGTGTGTGCGAA). The starting library was designed with a A:C:G:T molar ratio of 3:3:2:2.4 to adjust for equimolar amounts of nucleotide incorporation and primers. **Supplementary Table 1** summarizes the primers used in these studies. Dynabeads M-270 Carboxylic Acid were purchased from Thermo Fisher Scientific (Waltham, MA), along with N-hydroxysuccinimide (NHS) and 1-ethyl-3-(3-dimethylaminopropyl) carbodiimide hydrochloride required for magnetic bead activation. All RNA work was performed in designated areas following standard RNA workflow guidelines. All materials used were RNase-free (pipet tips, reagents, conical tubes, microcentrifuge tubes). All glassware was baked at 300°C for 2 hours in a standard oven. RNA workspaces were cleaned with RNase Zap (Thermo Fisher, AM9780). All buffers used were RNase-free. Binding buffer used throughout the selection was prepared with 20 mM Tris-HCl, 140 mM NaCl, 5 mM KCl, 1 mM MgCl_2_, and 1 mM CaCl_2_ (pH 7.4). Washing buffer was comprised of binding buffer supplemented with 0.005% Tween-20. Elution buffer was prepared with 50 mM Tris-HCl, 140 mM NaCl, and 50 mM EDTA (pH 7.4).

### Generation of the 2’-Fluorine Pyrimidine RNA Library

To generate the 2’-fluorine pyrimidine RNA aptamer library, 1 nmol of the starting ssDNA library was utilized. 60 standard Taq PCR reactions were prepared to amplify the starting ssDNA library in a total reaction volume of 3 mL (300 μL of Taq Buffer, 60 μL of dNTP, 180 μL of universal reverse primer, 180 μL of T7 forward primer, 10 μL of 100 μM N40 Library, 15 μL of standard Taq Polymerase, 2255 μL of ultrapure water). PCR was carried out according to the following cycling conditions: 95°C for 30 seconds, 8 cycles of [95°C for 30 seconds, 60°C for 60 seconds, 68°C for 60 seconds], and 68°C for 5 minutes. PCR reactions were pooled, and the resulting double-stranded DNA (dsDNA) was phenolchloroform extracted and ethanol precipitated. The dsDNA pellet was resuspended in 100 μL of ultrapure water, concentration was measured with Qubit Fluorometer (Thermo Fisher, Q33238), and correct band size (103 bp) was confirmed with gel electrophoresis on a 3% agarose gel. The dsDNA library was then converted to modified-base RNA using the Durascribe T7 Transcription Kit (Lucigen, MA170E). This kit produces RNA that is resistant to RNase A degradation through the replacement of canonical CTP and UTP with 2’-fluorine-dCTP (2’-F-dCTP) and 2’-fluorine-dUTP (2’-F-dUTP). The Durascribe kit uses a mutant T7 RNA polymerase that is able to efficiently incorporate 2’-F-dCTP, 2’-F-dUTP, ATP, and GTP into RNA transcription. 1 μg of dsDNA template was loaded into each IVT reaction. 10 transcription reactions were performed, with 20 μL of reaction buffer, 20 μL of ATP, 20 μL of 2’-F-dCTP, 20 μL of 2’-F-dUTP, 20 μL of GTP, 20 μL of DTT, and 20 μL of T7 Enzyme solution used in each reaction; an appropriate volume of dsDNA template was added based on its concentration and ultrapure water was then added to bring the total volume to 200 μL. Reactions were incubated at 37°C overnight, followed by incubation with 20 μL of DNase I for 15 minutes. RNA product was ethanol precipitated and resuspended in RNase-free AF Buffer.

### SELEX Workflow

In round 1 of selection, 1×10^4^ M-270 Carboxylic Acid Dynabeads were conjugated with 1 μg of human serum albumin and 1 μg of mouse serum albumin prior to incubation with the aptamer library. The magnetic beads were washed with MES buffer (25 mM 2-(N-morpholino)-ethane sulfonic acid, pH 6.0), activated with EDC/NHS chemistry for 30 minutes, and incubated with protein for 1.5 hours with rotation at room temperature. After incubation, the beads were washed with 50 mM Tris-HCl (pH 7.4) buffer and incubated with buffer for 1 hour to ensure that all unreacted groups on the magnetic beads were quenched. The beads were finally washed with PBS buffer with 0.005% Tween-20 and suspended in binding buffer and stored at 4°C until utilized. Beads were prepared fresh for each round of selection, and immediately prior to their use, beads were washed three times with wash buffer and resuspended in binding buffer.

Similar to our previous report^26^, a single well of a hydrogel bonded, ultra-low attachment 96-well plate (Corning, St. Louis, MO) was plumbed by drilling two holes into the lid, inserting dispensing needles (Jensen, North Andover, MA), and circulating fluid using a peristaltic pump (Fisher Scientific). Manifold pump tubing (PVC, 0.51 mm ID, Fisher Scientific) was flushed with washing buffer containing 100 mg/mL of yeast tRNA (Thermo Fisher Scientific) to passivate the lines before introducing the aptamer pools. Beads were trapped at the bottom of the plumbed well using a neodymium magnet. The selection was initiated with 5 nanomoles of starting RNA library. The starting library pool was heated to 95°C for 5 minutes and slowly cooled down to 25°C in a standard thermocycler at a rate of 0.5°C/min. Aptamer pools were circulated over the magnetically trapped beads at a rate of 20 mL/h. RNase-free wash buffer was then circulated at a rate of 50 mL/hour to continuously remove unbound and weakly bound aptamers. After washing, beads were resuspended in 100 μL of elution buffer and heated at 95°C for 10 minutes to elute the bound nucleic acids. Round 1 only included a positive incubation step with human and mouse albumin.

After elution, RNA was ethanol precipitated and resuspended in 10 μL of ultrapure water. Sunscript Reverse Transcriptase RNaseH (Expedeon 422050) was used to convert the eluted RNA to complementary DNA (cDNA). A reaction mixture was prepared with 10 μL of resuspended RNA, 2 μL of 10 μM universal reverse primer, 4 μL of 5X reaction buffer, 2 μL of 0.1 M DTT, 1 μL of 10 mM dNTP, 1.5 μL of Sunscript RT Enzyme. The reaction was run on a thermocycler using the following conditions: 65°C for 10 minutes, 70°C for 10 minutes, 75°C for 40 minutes, 95°C for 10 minutes. The cDNA product was PCR amplified with emulsion PCR using the T7 forward primer and universal reverse primer. dsDNA product was run on a 2% agarose gel and extracted with the Qiaex II Gel Extraction Kit (Qiagen, 20021). dsDNA was transcribed to RNA using 5 reactions of the Durascribe transcription kit as described above, heated at 37°C overnight, incubated with DNase I at 37°C for 15 minutes, and precipitated with ethanol. RNA product was then resuspended in AF Buffer and prepared for the next round of selection.

Round 2 and beyond included both positive incubation and negative counterselection steps. In the counterselection step, aptamer pools were circulated over quenched Dynabeads to remove aptamers binding to the surface of the beads. In these later rounds, we also decreased the time and number of protein-conjugated beads in the positive selection, while increasing the number of quenched beads and time of washing and counterselection. The full details of each round are found in **Supplementary Table 2**.

At the end of 5 rounds of selection, a TOPO-TA cloning kit (Thermo Fisher, 460572) was used to isolate individual aptamer sequences. The cloning reaction was set up with 4 μL of PCR product from the round 5 pool, 1 μL of salt solution, and 1 μL of TOPO vector. The reaction was gently mixed, incubated for 5 minutes at room temperature, and then placed on ice. One Shot Top10 Competent Cells were thawed on ice, mixed with 2 μL of the TOPO cloning reaction, and incubated on ice for 30 minutes. Cells were then heat-shocked for 30 seconds at 42°C, placed on ice, and mixed with 250 μL of SOC medium. The cells were rotated at 37°C for 1 hour, followed by spreading onto a pre-warmed ampicillin antibiotic selective plate, and incubated at 37°C overnight. The following day, individual colonies were picked and incubated overnight in 5 mL of LB broth with 50 μg/mL of ampicillin. dsDNA was extracted using the Plasmid Miniprep Kit (Qiagen, 27104). dsDNA concentration was measured using a Nanodrop (Fisher Scientific, ND-2000) and sequences were sent to Genewiz for Sanger Sequencing. Sanger Sequencing files were analyzed with SnapGene.

### Screening Aptamer Candidates

Nine prospective aptamers (**Table 1**) were ordered from IDT as ssDNA. These ssDNA aptamer sequences were PCR amplified into dsDNA with a T7 forward primer and unmodified reverse primer as described above. dsDNA aptamers were transcribed into RNA as described above. To screen RNA aptamer candidates for binding to Cy5-labeled human serum albumin, a Monolith Microscale Thermophoresis (MST) system was used (Nanotemper NT. 115). Briefly, aptamers were prepared by heating the mixture to 95°C for 5 minutes, snap-cooling on ice, and incubating at room temperature for 10 minutes. 40 nM solutions of Cy5-labeled albumin were prepared by diluting the protein in binding buffer. The Cy5-labeled albumin was mixed with each aptamer sample and incubated in the dark for 1 hour at room temperature. A standard Monolith NT.115 capillary was dipped into each solution and fluorescent dose-response was measured at 20% MST excitation power.

**Table 1.**
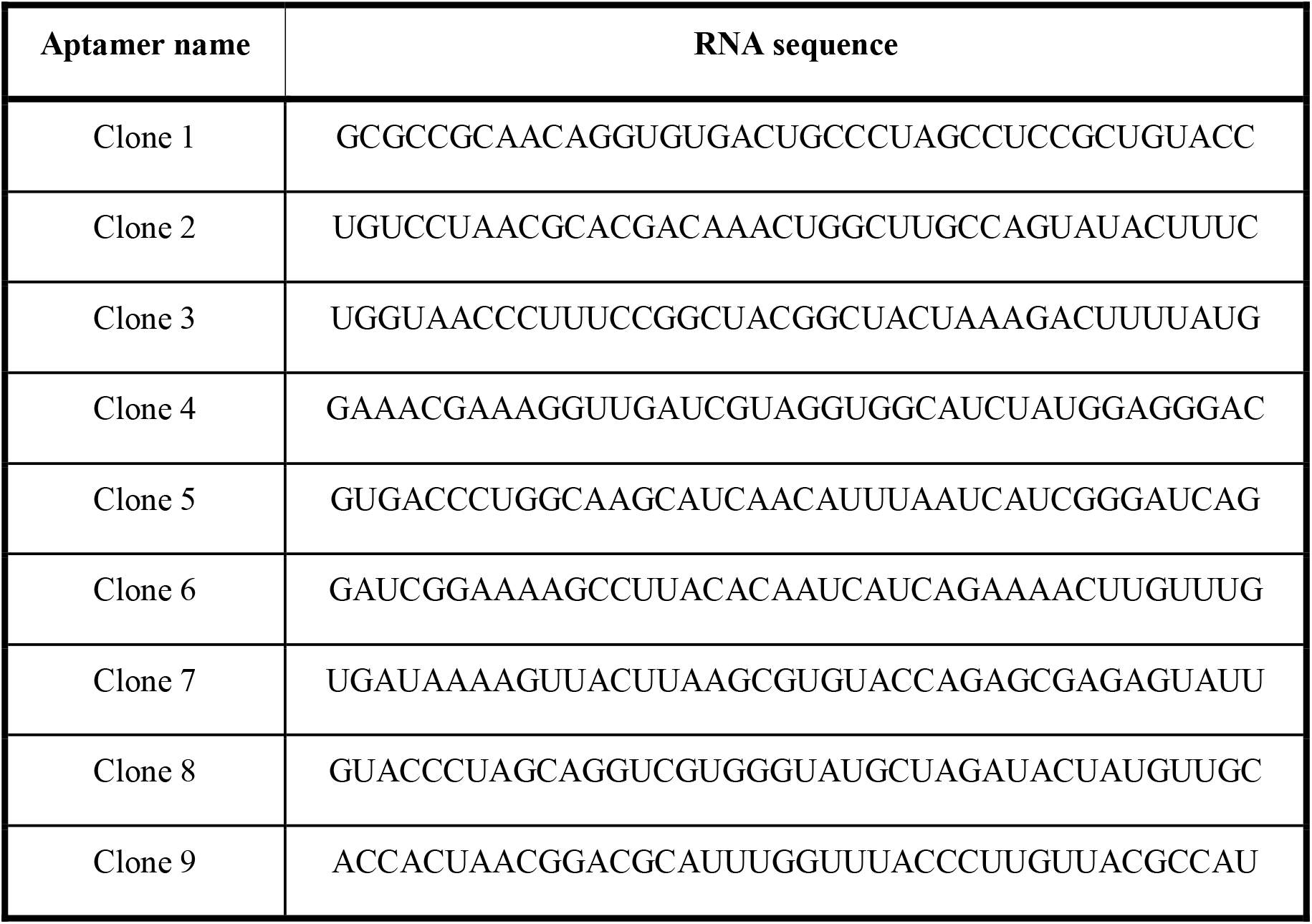
Prospective albumin-binding aptamers isolated by SELEX.

### Synthesis of Aptamer-siRNA Chimeras

Chimeras were generated by transcribing the RNA aptamer with the antisense strand of the siRNA separated by a two uracil base linker^15^. ssDNA encoding aptamer, the uracil base linker, and the antisense strand were ordered from IDT. The aptamer module was an albumin-binding clone or a scrambled control, and the siRNA module was either targeted to the luciferase gene or no gene (scrambled control). After PCR amplification and *in vitro* transcription of the aptamer-siRNA chimeras, the antisense strands were annealed to sense strands that were synthesized on a MerMade 12 Oligonucleotide synthesizer (BioAutomation). In brief, amidites were dissolved at 0.1 M in anhydrous acetonitrile and oligonucleotides were synthesized using standard coupling conditions. Mass fidelity was confirmed by LC ESI on a Waters Synapt. Depending on the experiment, the sense strands were either conjugated to Cy5 or left unconjugated. Aptamer antisense strands were annealed to sense strands by heating to 95°C for 5 minutes and slow cooling to room temperature for 1 hour. Annealing of aptamer antisense strand to sense strand was confirmed on a 3% agarose gel, comparing nucleic acid migration between the annealed chimeras and sense strand alone. Annealing of Cy5-labeled sense strand was similarly checked by visualizing nucleic acid migration by UV and IVIS Lumina III imaging (Caliper Life Science, Hopkinton, MA) (**Supplementary Figure 1**). After confirmation of annealing, the finalized aptamer-siRNA chimeras were used for downstream experiments. All chimera and siRNA sequences are shown in **Supplementary Table 3**.

### Affinity Measurements of Aptamer-siRNA Chimeras

The affinities of the Clone 1 and Clone 3 aptamer-siRNA chimeras for various targets were measured with a plate-based assay. Briefly, maleic anhydride amine-binding wells (Thermo Fisher, 15100) were washed three times with PBS containing 0.05% Tween-20. Equal amounts of mouse and human albumin were dissolved in immobilization buffer (PBS, pH 8.8) at a total concentration of 10 μg/mL. 100 μL of protein solution was incubated in each amine-binding well at 37°C for 1 hour. Protein solution was then removed and 200 μL of Protein Blocking Buffer (Thermo Fisher, 37515) was incubated in each well for 1 hour at room temperature. Blocking buffer was removed and the wells were washed three times with wash buffer. 2-fold serial dilutions of Cy5-labeled aptamer chimeras were prepared from 2 μM to 0 μM. Chimera dilutions were incubated in albumin-conjugated wells for 2 hours, washed three times with AF washing buffer, and Cy5 fluorescence was measured on a Tecan Infinite M1000 Pro plate reader. Data were fitted with nonlinear regression in Graphpad Prism to a one site, specific binding model to determine K_d_.

Clone 1 and Clone 3 specificities were further characterized for BSA/MSA/HSA versus IgG. In these experiments, the immobilization procedure described above was repeated for each moiety and 1 μM of chimera was incubated in each well for 2 hours, followed by washing and measurement of Cy5 fluorescence.

### Serum Stability Assessments

To determine the *in vitro* serum stability of Clone 1 and Clone 3 as well as a previously reported DNA aptamer (GTCTCAGCTACCTTACCGTATGTGGCCCAAAGCGTCTGGATGGCTATGAA) against human serum albumin^27^, 1 μg of each was incubated with 60% FBS at 37°C with agitation for the following time points: 0, 2, 4, 6, 8, and 24 hours. Each mixture was then run on a 3% agarose gel and visualized on an Odyssey Fc Imager after post staining with Gel Red.

### In Vitro Gene Knockdown Potency of Aptamer-siRNA Chimeras

Luciferase-expressing MDA-MB-231 cells were seeded overnight into a black 96-well plate at a density of 4,000 cells/well. 25 nM of aptamer-siRNA chimeras were complexed with Lipofectamine 2000 reagent (Thermo Fisher, L3000001) in Opti-MEM Media (Thermo Fisher, 31985062) according to the manufacturer’s protocol. Complexes were incubated with cells for 24 hours in normal culture media. Following the 24-hour incubation, treatments were removed and either replaced with regular culture media or luciferin-containing media (150 μg/mL) (Sigma, L9504) for evaluation of luminescence by IVIS imaging.

### Aptamer-siRNA Chimera Uptake in HUVECs

For imaging uptake of Cy5-labeled chimeras, HUVECs were seeded at a density of 15,000 cells/slide onto Lab-Tek II 8-well chamber slides and allowed to adhere overnight in a standard incubator. 150 nM aptamer-siRNA chimera sense and anti-sense strands were annealed for 5 minutes at 95°C followed by a 1-hour room temperature incubation. Cy5-labeled aptamer-chimeras were pre-complexed with HSA (Abcam, ab8030) at a 1:1 molar ratio for 30 minutes at room temperature and then added to the HUVECs in serum-free Opti-MEM media. After a 4-hour incubation, cells were washed with PBS, fixed in 4% PFA, and stained with DAPI. Imaging was performed on a Nikon Eclipse Ti-0E inverted microscopy base and images were analyzed using ImageJ software.

For quantifying uptake of Cy5-labeled chimeras, HUVECs were seeded at 25,000 cells/well in a 24-well plate and reached ~70% confluency after 24 hours. At this time point, HUVECs were treated with annealed Cy5-labeled aptamer chimeras (150 nM) for 2 hours in serum-free Opti-Mem. The aptamers were pre-complexed with HSA (Sigma, A7223) for 30 minutes at a 1:10 aptamer to HSA molar ratio prior to treatment. Cells were then prepared for flow cytometry by washing twice with PBS, harvesting with trypsin, and resuspending in PBS with 5% donkey serum. Mean fluorescence intensity was measured on a Guava EasyCyte (Luminex) after gating over 500 cellular events. **Supplemental Figure 2** shows the gating strategy used for analysis.

### Intravital Microscopy

Male CD1 mice aged 4-6 weeks old (Charles River Laboratories) were anesthetized using isoflurane and immobilized on a heated confocal microscope stage. Mouse ears were cleaned with a depilatory cream, and microscope immersion fluid was using to immobilize the ear on a glass coverslip. An ear vein was detected using a light microscope and brought into focus such that the flowing red blood cells were clearly visible. Once focused, the microscope (Nikon C1si+) was switched to confocal laser mode and set to acquire images continuously every second. The mouse was then injected *via* tail vein with 2 nmol of Cy5-labeled aptamer-siRNA chimera or relevant controls (approximately 1 mg/kg), and images were collected continuously for approximately 30 minutes. Fluorescence was evaluated by averaging pixel intensities within the circular region of interest located within the ear vein in focus. Maximum initial fluorescence in the vein was set to a time of 0 seconds, and data were fit to a one-compartment model in PKSolver to determine pharmacokinetic parameters by examining the data collected from 0 to 20 minutes.

### Statistics

Groups were statistically compared using a one-way ANOVA test (unpaired) or two-way ANOVA (paired) test with Tukey’s or Dunnett’s comparisons. For comparison between two groups, a Mann Whitney test was used. A p-value <0.05 was deemed representative of a significant difference between groups. For all data shown, the arithmetic mean and standard deviation are reported, and the sample size (n) is indicated.

### Ethics Statement

The animal studies described were conducted with adherence to the *Guide for the Care and Use of Laboratory Animals*. All experiments with animals were approved by Vanderbilt University’s Institute for Animal Care and Use Committee.

## Results

### Choice of Target and Outline of SELEX Process

To identify a high affinity and specificity aptamer with prospective preclinical and clinical relevance, five rounds of SELEX were completed with both human and mouse serum albumin as the primary targets and quenched micromagnetic beads as the negative counterselection targets. In the first round of selection, only a positive selection with the human and mouse albumin was performed. In rounds 2 to 5, a negative selection step was performed immediately after positive selection with the quenched micromagnetic beads. With each consecutive round, the amount of primary target and the time of positive selection were decreased, while the amount of off-target, the time of washing, and the time of negative selection were increased (**Supplementary Table 2**); this approach increases the selection pressure through each round of selection and helps select aptamers with higher affinity for their target^28, 29^. After the fifth round of selection, the nucleic acid pool was cloned into a bacterial plasmid, and individual colonies were isolated and sequenced to identify candidate aptamers (**Table 1**).

### Characterization of Prospective Aptamers

Nine aptamer candidates generated from SELEX were screened for their binding to Cy5-labeled human serum albumin using microscale thermophoresis (MST) (**Figure 1A**). MST measures two parameters: response amplitude, which is the signal difference between the bound and unbound state of the fluorescent molecule, and signal to noise ratio, which is the response amplitude divided by measurement noise (average standard deviations of all points from the fitted curve). A true binding signal should have a response amplitude and a signal to noise ratio of at least 5 response units. Clone 3 had the highest response amplitude of 10.8 ± 2.2 fluorescent units and the highest signal to noise ratio of 10.1 ± 1.2, while Clone 1 had the second highest response amplitude of 8.4 ± 1.8 fluorescent units and an acceptable signal to noise ratio of 5.2 ± 2.0. These values for the other clones were either borderline or did not meet the required threshold for both parameters. Thus, Clone 1 and Clone 3 were selected for further characterization.

**Figure 1.**
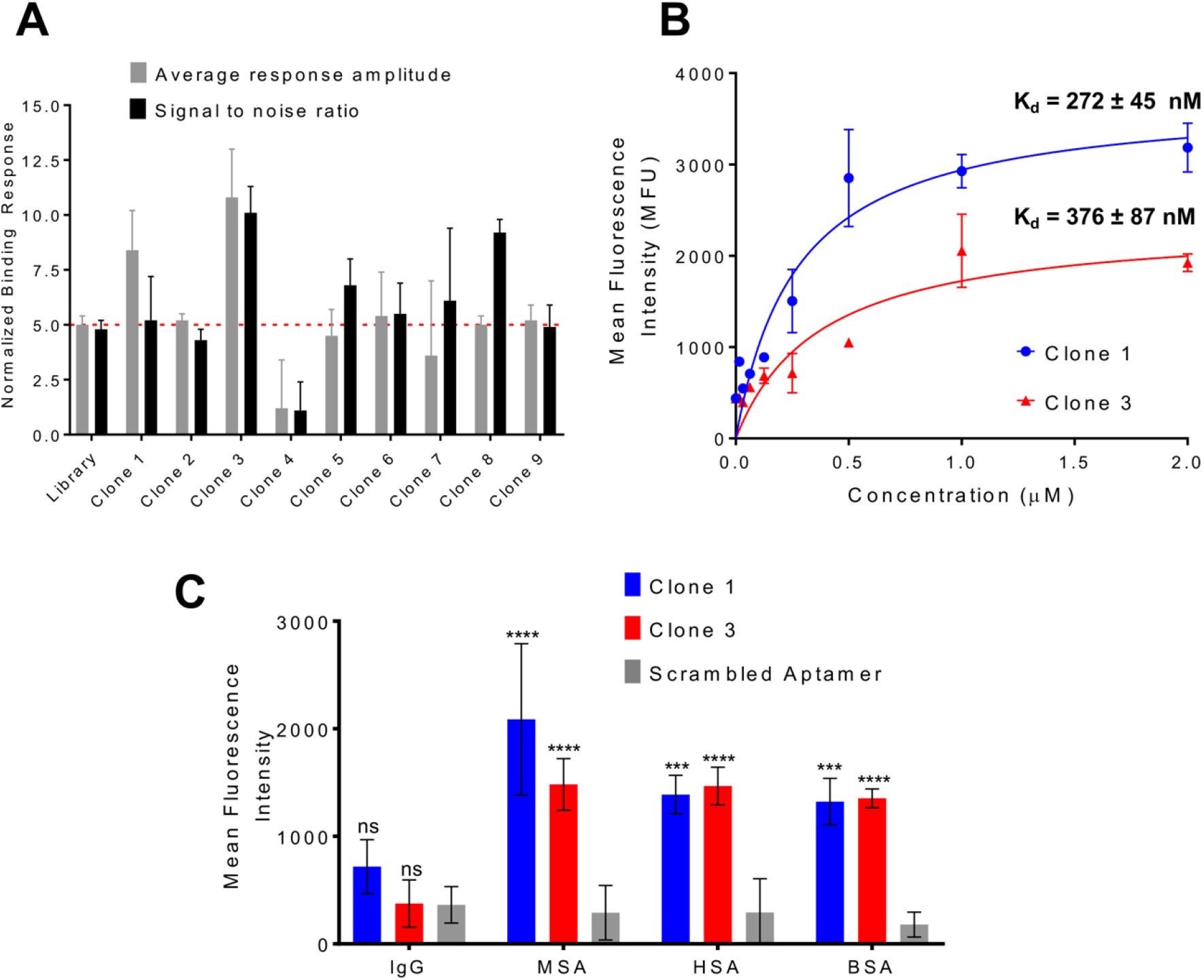
Characterization of albumin-binding aptamer affinity and specificity. **A)** Aptamer candidates identified via SELEX were screened by microscale thermophoresis for their affinity for human serum albumin (n=3). Clones that demonstrated an average response amplitude and signal to noise ratio above 5 are selected for further investigation. **B)** Clones 1 and 3 were formulated into aptamer-siRNA chimeras and affinity was measured against a mixture of immobilized mouse and human albumin. Data were fitted to a one site binding model to calculate dissociation constants (K_d_) (n=6). **C)** Relative binding of Clone 1, Clone 3, and a scrambled aptamer control to immobilized mouse albumin (MSA), human albumin (HSA), bovine albumin (BSA), and IgG (n=3). Statistical significance was assessed with a twoway ANOVA with Dunnett’s multiple comparison’s test (****, p<0.0001; ***, p<0.001).

Chimeric aptamer-siRNAs were generated using a two nucleobase uracil (UU) linkage between the aptamer and the antisense strand of a Dicer substrate RNA. This design builds on previously reports that support the use of this linker in connecting aptamers to Dicer substrate RNAs, and that appendage to the antisense strand is preferable to the sense strand for silencing potency^15^. The affinities of Clone 1 and Clone 3 chimeras for albumin were explicitly measured for binding to immobilized, unlabeled human and mouse serum albumin. For these measurements, equal amounts of human and mouse serum albumin were conjugated to the surface of maleic anhydride-coated wells and incubated with varying concentrations of fluorescent chimera – in this case, the sense strand annealed to the chimera was labeled with Cy5. After incubation and washing, the amount of bound aptamer relative to the concentration added to each well can be used to extrapolate a binding affinity. Here, Clone 1 exhibited a K_d_ of 272 ± 45 nM and Clone 3 exhibited a K_d_ of 376 ± 87 nM (**Figure 1B**). We also tested binding of Clone 1, Clone 3, and scrambled chimeras for binding to albumin from various species compared to IgG to assess their specificity. Here, the chimeras selected for albumin binding demonstrated similar binding as a scrambled chimera for IgG, but several orders of magnitude greater binding for different species of albumin (**Figure 1C**). To build on these results, we further explored whether the sequences of Clone 1 and Clone 3 could be truncated while retaining affinity for albumin (**Figure 2A, B)**^30^. However, in all cases the truncated aptamers (**Figure 2C**) exhibited negligible binding relative to a negative control and empty well (**Figure 2D**). Overall, our results demonstrate that Clone 1 and Clone 3, in both their unmodified aptamer and aptamer-chimera forms, bind albumin with high affinity and specificity in a sequence-specific manner.

**Figure 2.**
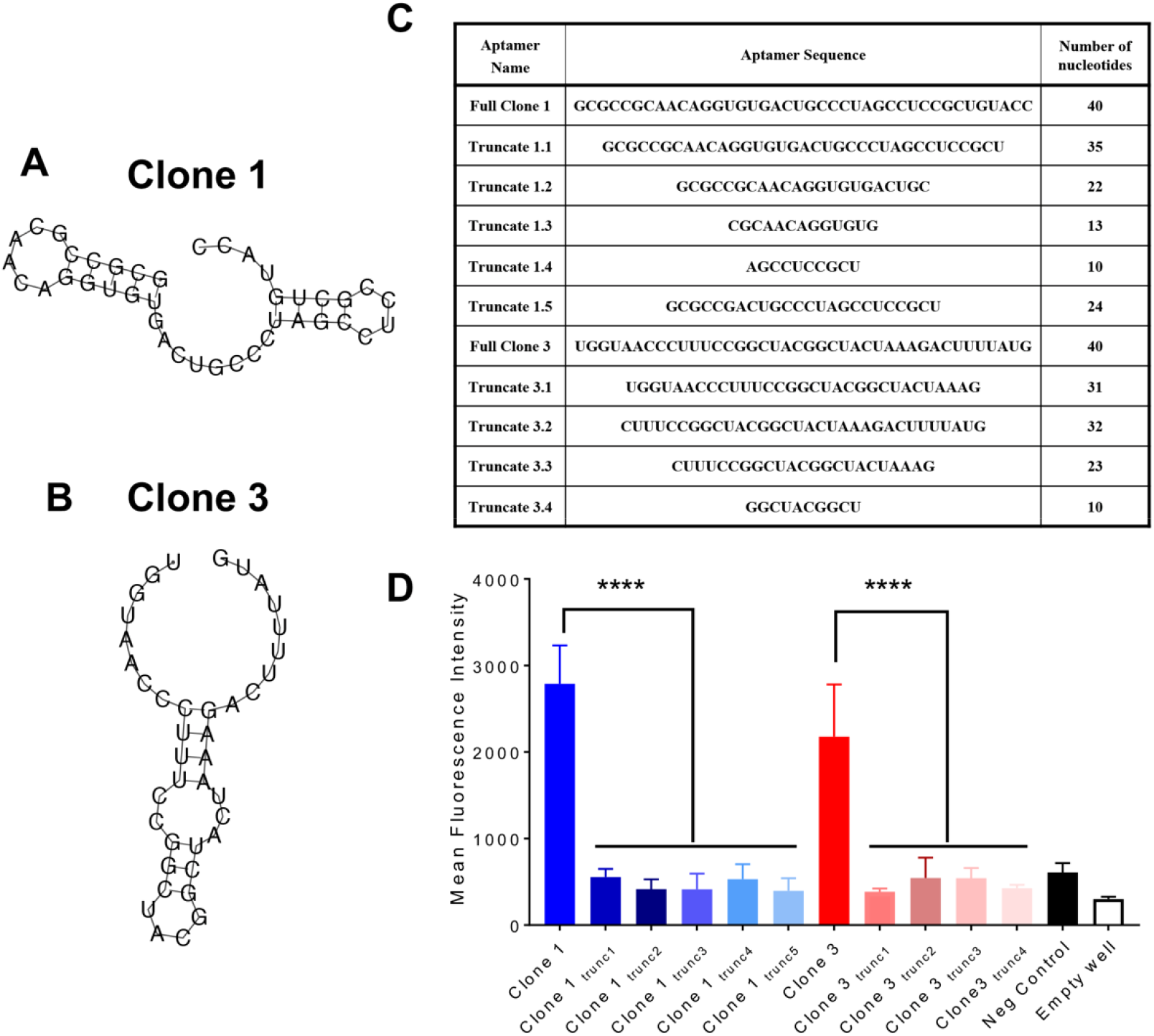
Full aptamer sequence is required for albumin binding. **A)** Predicted structure of Clone 1 from the Vienna RNA Websuite. **B)** Predicted structure of clone 3 from the Vienna RNA Websuite. **C)** Truncation sequences for Clone 1 and Clone 3 aptamers. **D)** Relative binding of Clone 1, Clone 3, and truncated variants to immobilized mouse and human albumin (n=3). Statistical significance was assessed with an ordinary one-way ANOVA with Sidak’s multiple comparison test (****, p<0.0001).

### In Vitro Characterization of Aptamer-siRNA Chimera Cellular Uptake and Silencing Potency

To examine the effect of albumin affinity/binding of our chimeras on cell uptake, we treated HUVECs, an endothelial cell line, with our candidate chimeras that had been precomplexed with albumin. Importantly, endothelial cells are known for their uptake and transport of albumin *in vivo*^31^. Flow cytometry demonstrated significantly higher uptake of albumin-binding Clone 1 versus a scrambled aptamer-chimera control when the aptamers were pre-mixed with albumin prior to application onto HUVECs (**Figure 3A**). Imaging of adhered HUVECs further demonstrated qualitatively improved uptake of the albumin-binding chimeras, as visualized by small puncta (**Figure 3B**).

**Figure 3.**
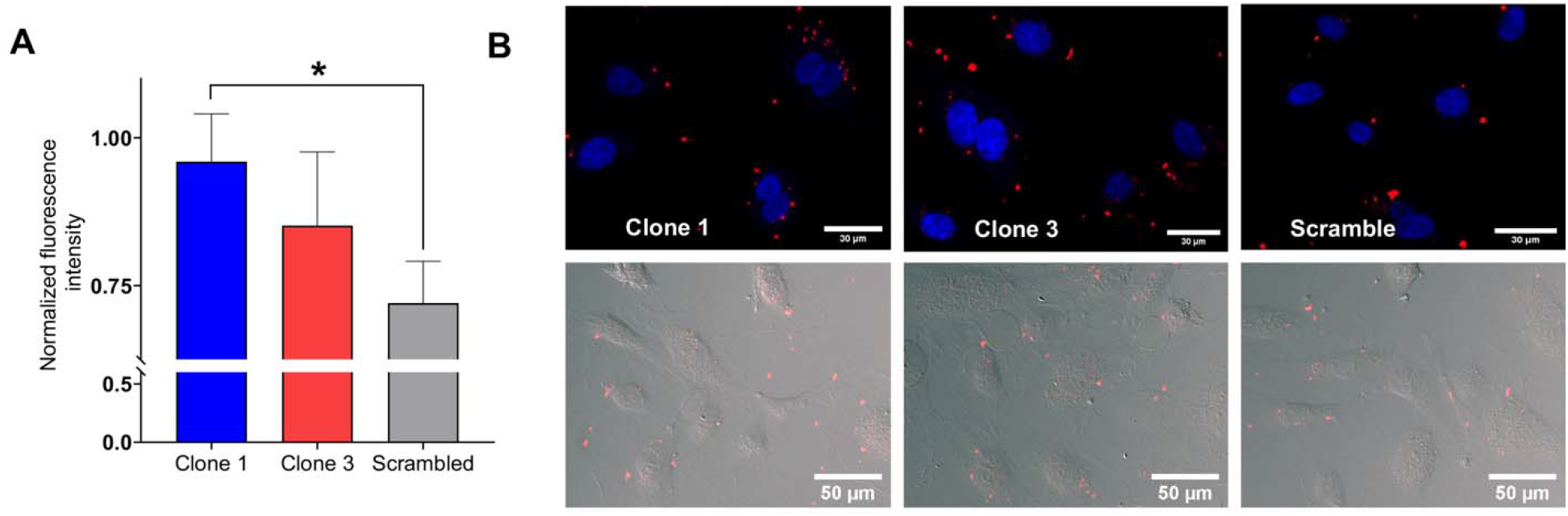
Uptake of aptamer chimeras in HUVECs. **A)** Quantitative assessment of aptamer chimera uptake by flow cytometry (n=4). Mean cell fluorescence intensities are normalized to the highest intensity within each biological replicate. Statistical significance was assessed by a one-way ANOVA with Tukey’s multiple comparison test (*, p<0.05). **B)** Qualitative assessment of aptamer chimera uptake by fluorescence microscopy.

To further validate our chimera constructs, we sought to examine the *in vitro* knockdown capability of our candidates with different designs. Luciferase-expressing MDA-MB-231 cells were used to compare the knockdown efficiency of the chimeras with their unappended counterparts (i.e. free siRNA). The proceeding experiments used chimeras with the antisense strand appended to the aptamer and annealed to the sense strand. The fusion of the siRNA with the aptamer did not produce any significant differences in silencing 48 hours after delivery *via* lipofection (**Figure 4A**). We sought to bolster this finding by additionally testing the aptamer chimera fused to siRNA targeting Luciferase versus a negative control siRNA (**Figure 4B**). We determined that the magnitude of knockdown was consistent between the negative control free siRNA and the same negative control siRNA fused to the aptamers (**Figure 4B** compared to **Figure 4A**). Overall, the aptamer-siRNA chimeras are able to retain their silencing potency *in vitro* with about 50% knockdown efficiency compared to the scrambled controls.

**Figure 4.**
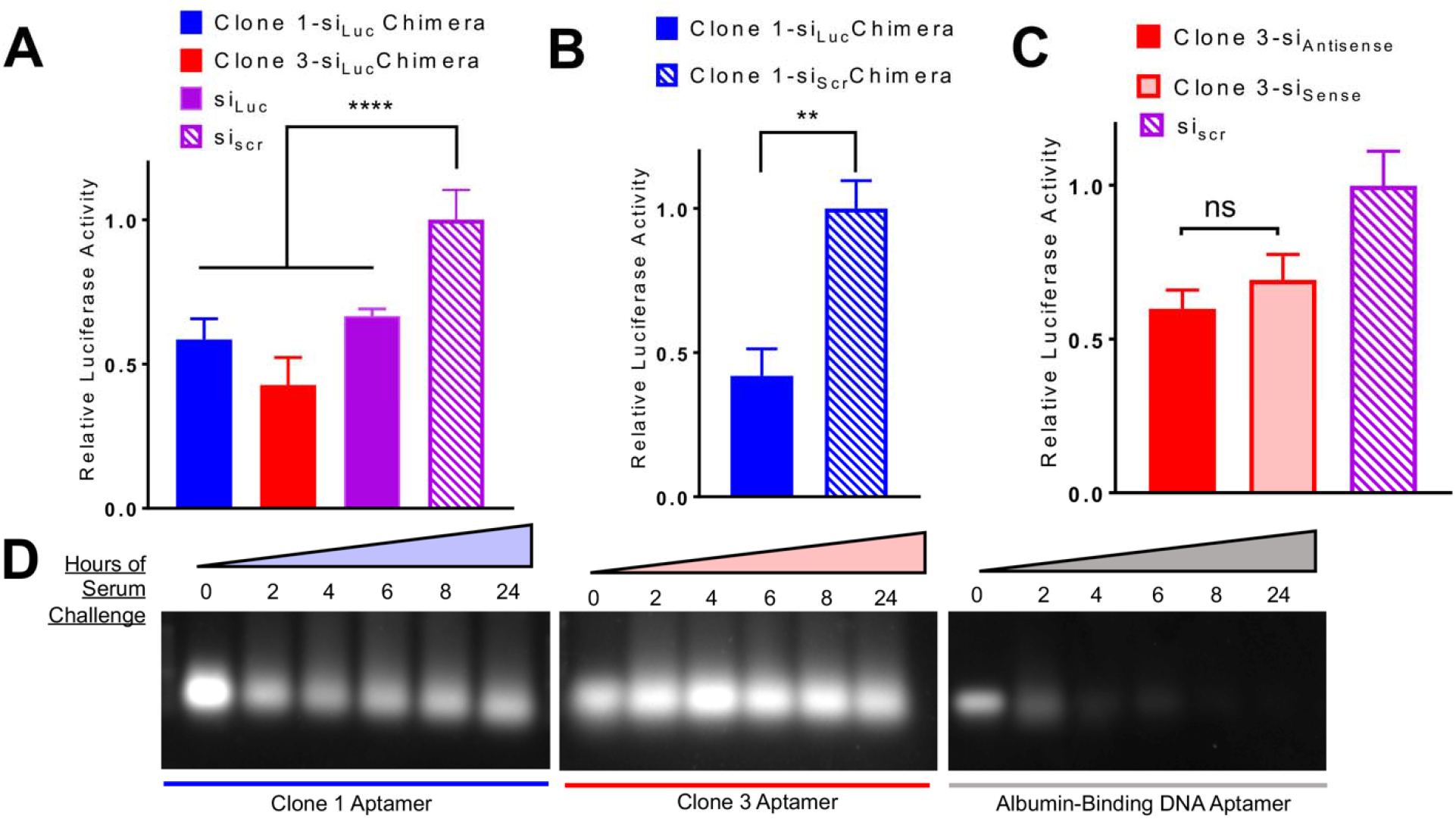
Assessment of aptamer chimera knockdown potency and serum stability. **A)** Silencing potency of Luc-targeting Clone 1 and Clone 3 aptamer chimeras relative to free Luciferase-targeting siRNA and scrambled siRNA control in MDA-MB-231 cells (n=6). Statistical significance was assessed using a one-way ANOVA with Dunnett’s multiple comparisons test (****, p<0.0001). **B)** Silencing potency of Luciferase-targeting Clone 1 aptamer chimera versus Clone 1 aptamer chimera harboring a scrambled siRNA in MDA-MB-231 cells (n=6). Statistical significance was assessed by a Mann-Whitney test (**, p<0.01). **C)** Silencing potency of Luciferase-targeting Clone 3 aptamer chimeras in sense versus antisense orientation relative to a scrambled siRNA control in MDA-MB-231 cells (n=3). Statistical significance was assessed with a one-way ANOVA with Tukey’s multiple comparisons test. **D)** Serum stability of Clone 1 and Clone 3 aptamers relative to a previously published albumin-binding DNA aptamer.

Aptamer-siRNA chimeras can be designed with either the siRNA antisense strand or sense strand appended to the aptamer. We additionally sought to confirm that appendage to the antisense maintained target knockdown potency. Appendage of our chimera to either the 5’ end of the sense or antisense strand of the Dicer substrate siRNA against Luciferase did not show appreciable differences in knockdown potency (**Figure 4C**).

RNA aptamers are generally preferred to DNA aptamers with regard to the diversity of binding conformations and subsequent increased likelihood of libraries containing sequences that are hits for a target. Additionally, the serum stability of 2’-fluorine-modified pyrimidine RNA aptamers is appealing because of their resistance to degradation by nucleases when administered systemically^32–34^. Since there is a previously reported DNA aptamer against human serum albumin^27^, we sought to compare the stability of our 2’F modified RNA chimeras to this DNA aptamer by challenging all constructs with 60% serum over 24 hours **(Figure 4D)**. Modified-base chimeras are able to resist degradation over the 24-hour incubation period whereas a DNA-based alternative could not, indicating the effectiveness of the 2’F bases in reducing aptamer degradation by nucleases.

### In Vivo Serum Half-Life of Aptamer-siRNA Chimeras

To test the ability of the albumin-binding aptamer-chimeras to extend the bioavailability of siRNA, we assessed circulation time using intravital microscopy. We continuously monitored Cy5 fluorescence of labeled aptamer-chimeras in mouse ear vasculature after intravenous administration (**Figure 5A**), which allowed us to measure fluorescence decay and subsequent pharmacokinetic parameters. The Clone 1 chimera showed a statistically significant 1.6-fold improvement in absolute circulation half-life (12.6 ± 0.93 minutes) compared to the scramble aptamer-chimera (7.46 ± 1.26 minutes) used as a negative control, whereas Clone 3 chimera showed a modest but insignificant improvement (7.94 ± 1.86 minutes) (**Figure 5B**). Our findings suggest that sequence-specific affinity of the Clone 1 aptamer for albumin yields improved bioavailability of the chimera.

**Figure 5.**
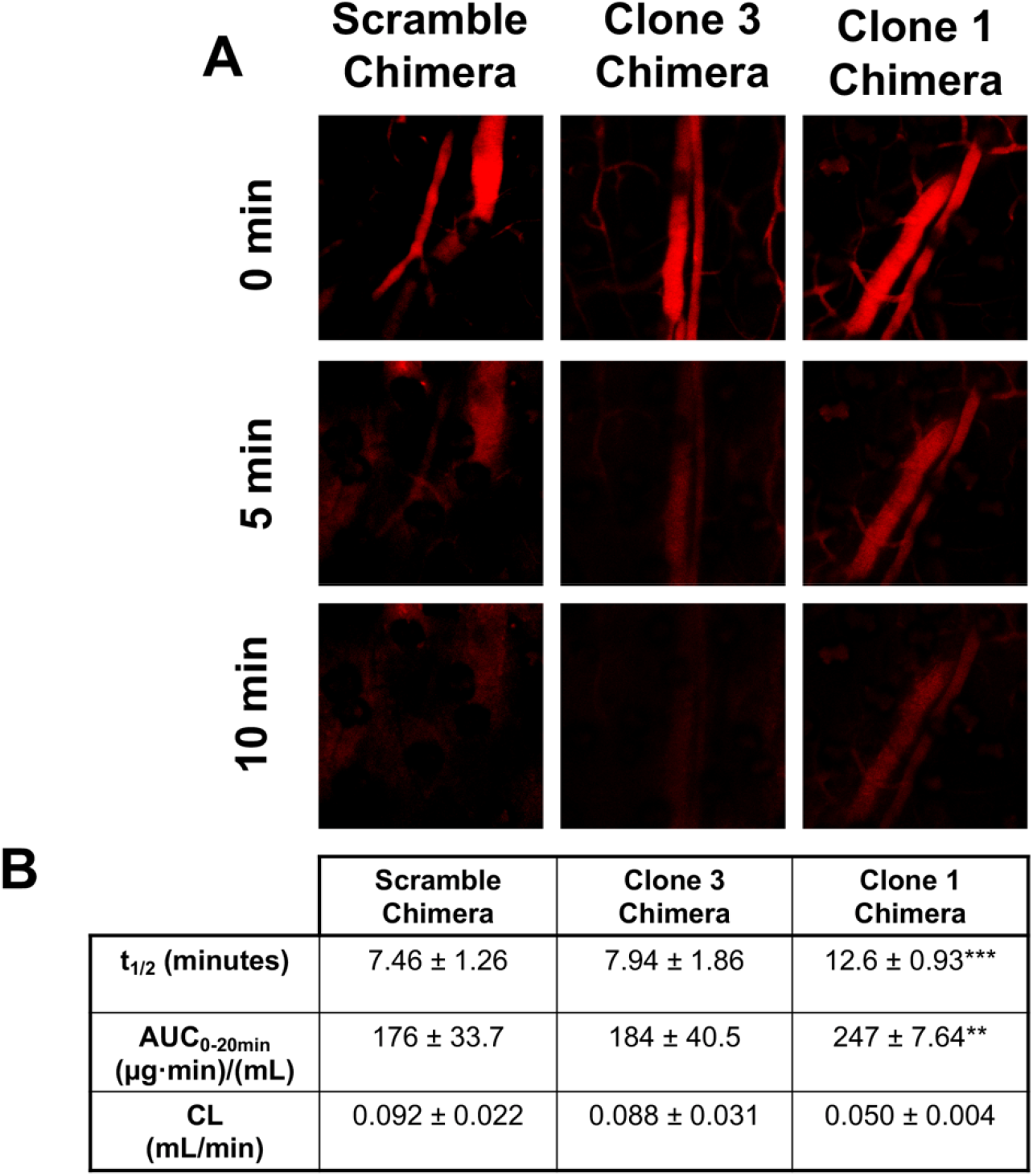
Aptamer chimera pharmacokinetic parameters calculated using intravital microscopy. **A)** Representative fluorescent images of mouse ear vasculature after intravenous injection of aptamer chimeras. **B)** Pharmacokinetic parameters calculated using PK Solver. Statistical significance was assessed with a two-way ANOVA with Tukey’s multiple comparisons test (***, p<0.001; **, p<0.01).

## Discussion

Herein, we describe the isolation of albumin-binding RNA aptamers, followed by the formulation and characterization of aptamer-siRNA chimeras. An optimized SELEX procedure was performed with modified RNA bases to generate serum-stable aptamers that bind albumin with high affinity. Two candidate aptamers (Clone 1 and Clone 3) were then fused to siRNA to form aptamer-siRNA chimeras, which exhibited potent gene silencing activity *in vitro*. The Clone 1 chimera, which possessed the highest affinity for albumin, further exhibited increased circulation half-life *in vivo*.

We suggest that the modular design of aptamer-siRNA chimeras could facilitate broad applications of gene-targeted therapeutics. While we employed a fused aptamer-siRNA design for these proof-of-concept studies, there are many programmable nucleic acid-based nanostructures that could incorporate albumin-binding aptamers to enhance *in vivo* half-life, such as self-assembled tetrahedral structures. Through complementary base pairing, these and other structures can be loaded with siRNA and decorated with multiple types of aptamers. In this scenario, the albumin-binding aptamers could prospectively increase circulation time, while other aptamers could be used to target specific cell types. For example, we recently described a cell-SELEX strategy to generate aptamers that can discriminate membrane receptor homologs on the surface of living cells^35^, which could be extended here to preferentially target and retain multifaceted nucleic acid nanostructures within different tissue compartments. This strategy could be particularly powerful in solid tumors, where we have shown preferential penetration and gene silencing of albumin-binding nanocomplexes^8^, which could potentially be augmented by using aptamers that bind receptors that are overexpressed or mutated on the surface of the cancer cells.

To achieve this goal, some improvements to the existing albumin-binding aptamer platform will be necessary. First, the aptamer-siRNA chimeras are produced by PCR amplification of ssDNA templates, followed by *in vitro* transcription to the final RNA construct. This approach does not produce high yields of RNA (~1 μg per transcription) and can generate truncated byproducts. Solid-phase synthesis would likely increase yield and product consistency, but synthesis efficiency is inversely proportional to length of the oligonucleotide. Our attempts to truncate the albumin-binding aptamer, which would have made solid-state synthesis viable, were not successful. For future work, the SELEX process likely needs to be repeated with a smaller 20-mer library such that the final aptamer-siRNA chimeras would be a manageable length for solid-state synthesis.

Second, it may be of interest to adjust the affinity and specificity of the aptamer for albumin, as well as other properties that could influence aptamer-siRNA chimera potency. Based on our intravital microscopy experiments, we presume that increasing the affinity of aptamers for albumin could further improve the circulation half-life of the aptamer-siRNA chimeras, but altering affinity could also influence biodistribution, cellular uptake, and endolysosomal escape. Although we provided evidence that the aptamer-siRNA chimeras maintain their *in vitro* gene knockdown potency when delivered *via* lipofection, they will still be hindered by their innate inability to escape the endolysosomal pathway when delivered alone. We suggest that further functionalization of the aptamer-siRNA chimeras with synthetic moieties that disrupt endosomes could enhance silencing activity. Such modifications could be built directly into the aptamer-siRNA chimeras or onto more complex nucleic acid nanostructures. Overall, our experimental evidence suggests that aptamer-siRNA chimeras represent an exciting new avenue for improving the bioavailability of siRNA, and future work will focus on making improvements to the platform to improve its translatability.

## Acknowledgments

Funding for this work was provided by the Chan Zuckerberg Initiative (grant 2018-191850 to ESL), the BrightFocus Foundation (grant A20170945 to ESL), and National Institutes of Health grants R01 CA224241 and R01 EB019409 (to CLD). ENH was supported by a National Science Foundation Graduate Research Fellowship.

## Author contributions

JCR, ENH, CLD, and ESL conceived the research plan. JCR, ENH, and AGS carried out experiments. All authors wrote, reviewed, and edited the manuscript.

## Notes

### Competing Interest Statement

The authors have declared no competing interest.

