## Supplemental information for "Albumin-binding Aptamer Chimeras for Improved siRNA Bioavailability"

**Supplementary Table 1. Primer sequences.**

| **Primer name** | **Primer sequence** |
| --- | --- |
| **T7 forward primer** | **GTATAATACGACTCACTATAGGG** |
| **Universal reverse primer** | **TTCGCACACACGGACTTACG** |
| **Luc Sense reverse primer** | **GAGGAGTTCATTATCAGTGCAATTG** |
| **Luc Antisense reverse primer** | **AACAATTGCACTGATAATGAACTCTC** |
| **Scrambled Sense reverse primer** | **ATACGCGTATTATACGCGATTAACG** |
| **Scrambled Antisense reverse primer** | **GTCGTTAATCGCGTATAATACGCGTAT** |

**Supplementary Table 2. Parameters for SELEX.**

| **Round** | **Amount of mouse and human albumin (ng)** | **Amount of quenched beads (mg)** | **Time of positive selection (min)** | **Time of washing and counterselection (min)** |
| --- | --- | --- | --- | --- |
| 1 | 2000 | - | 60 | - |
| 2 | 1000 | 1 | 50 | 30 |
| 3 | 500 | 2 | 45 | 45 |
| 4 | 100 | 5 | 40 | 60 |
| 5 | 100 | 10 | 30 | 60 |

**Supplementary Table 3. siRNA and chimera sequences.**

| **Designation** | **Sequence** | **Number of nucleotides** |
| --- | --- | --- |
| **Luc Sense** | **CAAUUGCACUGAUAAUGAACUCCTC** | **25** |
| **Luc Antisense** | **GAGGAGUUCAUUAUCAGUGCAAUUGUU** | **27** |
| **Scramble Sense** | **CGUUAAUCGCGUAUAAUACGCGUAU** | **25** |
| **Scramble Antisense** | **AUACGCGUAUUAUACGCGAUUAACGAC** | **27** |
| **Clone 1 Luc Antisense** | **GCGCCGCAACAGGUGUGACUGCCCUAGCCUCCGCUGUACCAAGAGGAGUUCAUUAUCAGUGCAAUUGUU** | **69** |
| **Clone 3 Luc Antisense** | **UGGUAACCCUUUCCGGCUACGGCUACUAAAGACUUUUAUGAAGAGGAGUUCAUUAUCAGUGCAAUUGUU** | **69** |
| **Negative Control Luc Antisense** | **GAUACUGAGCAUCGUACAUGAUCCCGCAACGGGCAGUAUUAAGAGGAGUUCAUUAUCAGUGCAAUUGU** | **69** |
| **Clone 1 Luc Scr Antisense** | **GCGCCGCAACAGGUGUGACUGCCCUAGCCUCCGCUGUACCAAAUACGCGUAUUAUACGCGAUUAACGAC** | **69** |
| **Clone 1 Luc Sense** | **GCGCCGCAACAGGUGUGACUGCCCUAGCCUCCGCUGUACCAACAAUUGCACUGAUAAUGAACUCCUC** | **67** |
| **Clone 3 Luc Sense** | **UGGUAACCCUUUCCGGCUACGGCUACUAAAGACUUUUAUGAACAAUUGCACUGAUAAUGAACUCCUC** | **67** |
| **Negative Control Luc Sense** | **GAUACUGAGCAUCGUACAUGAUCCCGCAACGGGCAGUAUUAACAAUUGCACUGAUAAUGAACUCUCUC** | **67** |
| **Clone 1 Scr Luc Sense** | **GCGCCGCAACAGGUGUGACUGCCCUAGCCUCCGCUGUACCAACGUUAAUCGCGUAUAAUACGCGUAU** | **67** |
| **Clone 3 Scr Luc Sense** | **UGGUAACCCUUUCCGGCUACGGCUACUAAAGACUUUUAUGAACGUUAAUCGCGUAUAAUACGCGUAU** | **67** |
| **Negative Control Scr Luc Sense** | **GAUACUGAGCAUCGUACAUGAUCCCGCAACGGGCAGUAUUAACGUUAAUCGCGUAUAAUACGCGUAU** | **67** |

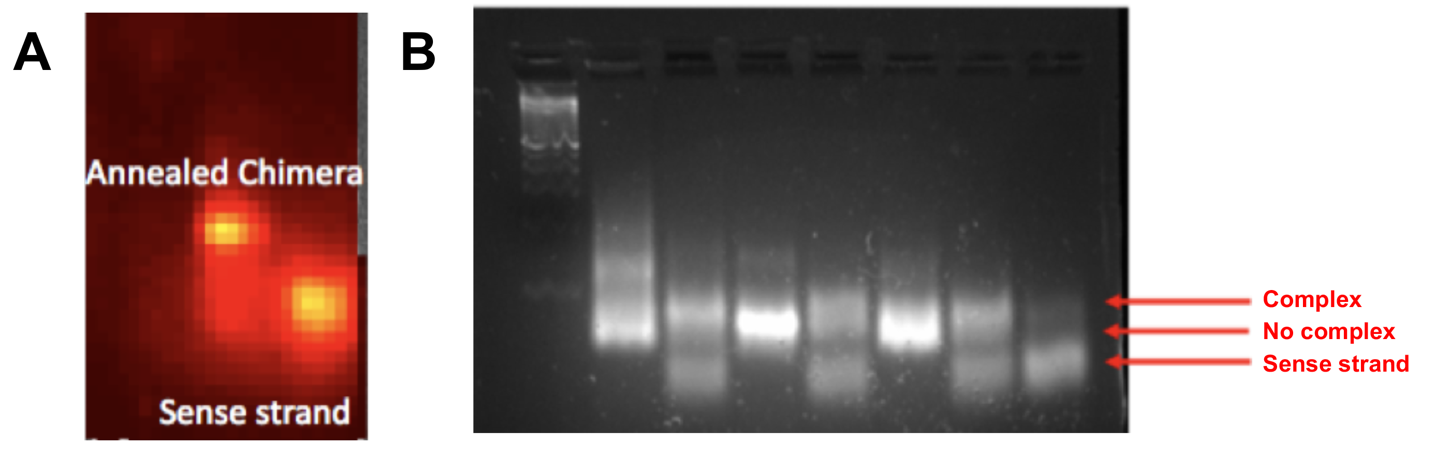

**Supplementary Figure 1. Confirmation of annealed chimeras on 3% agarose gels. A)** Fluorescently labeled sense strand was visualized on an IVIS system. **B)** UV imaging after Gel Red post-staining.

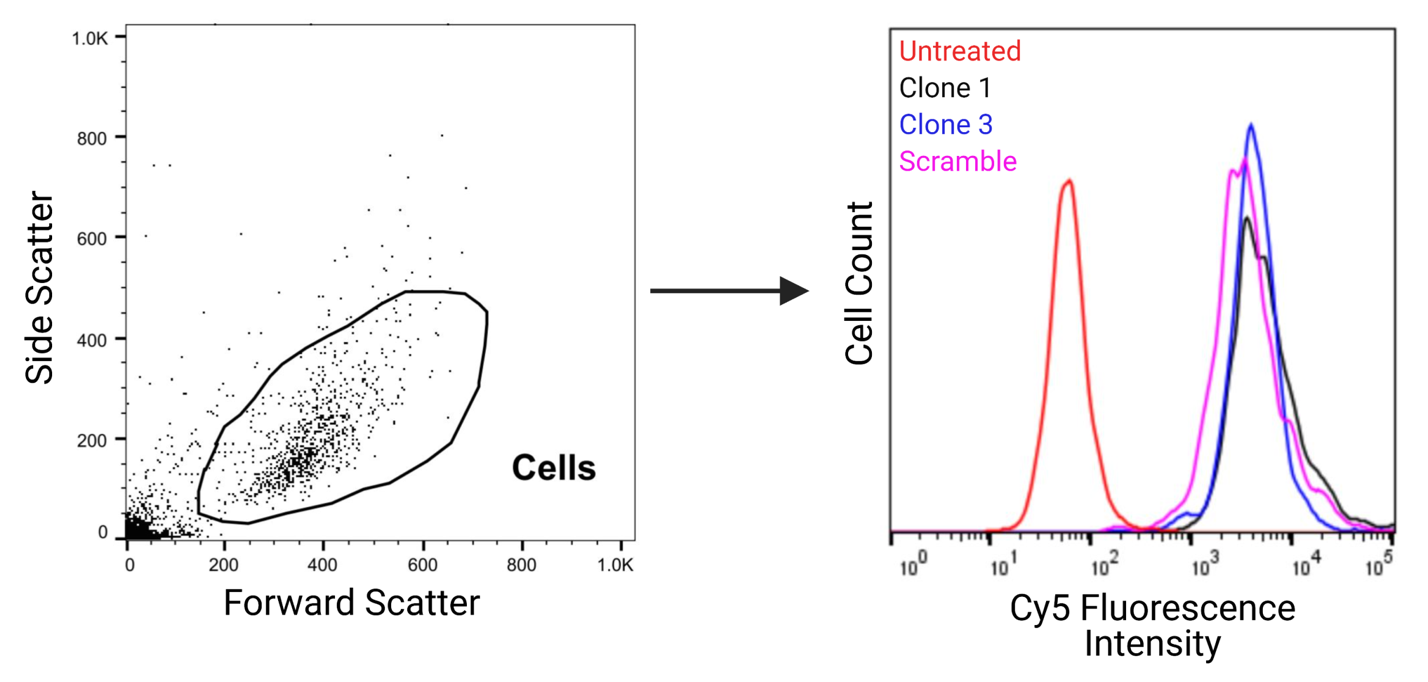

**Supplementary Figure 2. Gating strategy for assessing uptake of albumin-binding aptamers in HUVECs.**
